# Hydrogel Fiber Endomicroscopy

**DOI:** 10.64898/2026.03.23.713710

**Authors:** Peng Chen, Keyi Han, Zijun Gao, Claire M. Deng, Haoyu Xu, Zhi Ling, Corey Zheng, Mithila Sawant, Marcus T. Cicerone, Aparna H. Kesarwala, Jeffrey E. Markowitz, Shu Jia

## Abstract

Multimode fibers enable minimally invasive, high-resolution imaging through ultrathin probes, thereby enhancing diagnostic precision and facilitating real-time monitoring in delicate anatomical regions. In this work, we introduce HYFEN, a hydrogel-based endomicroscopic imaging platform for flexible, biocompatible, and subcellular-scale fluorescence microscopy. HYFEN leverages the unique properties of hydrogel materials, adaptive optics, and pixel-wise image enhancement to address challenges associated with silica-based fibers, including mode scrambling, limited field of view, and mechanical rigidity. The technique achieves precise mode threading, rapid diffraction-limited focusing at kilohertz speeds, and high-fidelity fluorescence signal acquisition with subcellular resolution. Notably, the approach extends fluorescence imaging under enhanced fiber dimensions and bending conditions that are unachievable with conventional modalities. Together, these advances establish HYFEN as a versatile platform for next-generation biointerfacing and minimally invasive imaging across biomedical and clinical settings.

## INTRODUCTION

Optical endoscopy has substantially advanced medical diagnostics by providing minimally invasive, high-resolution visualization of internal tissues^1-5^. Among major endoscopic modalities, multimode fiber (MMF)-based imaging has gained notable traction, exploiting a single ultrathin fiber capable of transmitting complex optical fields through thousands of guided modes, effectively replacing conventional, bulky fiber bundles^6,7^. Despite their advantages, MMFs are inherently hindered by optical and material constraints, including modal dispersion, mode scrambling, and sensitivity to fiber bending, which degrade the transmitted wavefront into a randomized speckle pattern that impairs image fidelity^8^. Recent advances in light-field computation and wavefront shaping have alleviated these limitations, enabling diffraction-limited focusing and accurate image reconstruction under controlled conditions^9,10^. Consequently, MMFs have been widely employed in biomedical imaging applications, including minimally invasive endoscopy, fluorescence microscopy, label-free optical tomography, and deep-tissue interrogation^11-20^. Nonetheless, conventional MMFs, typically fabricated from rigid silica or polymer materials, remain suboptimal for interfacing with flexible and deformable biological environments due to their mechanical stiffness and fragility^17,21^.

To overcome these limitations, optical fibers made from soft and biocompatible materials, such as silk^22^, cellulose^23^, and bacteria^24^, have been explored. Among these alternatives, hydrogel-based optical fibers have recently garnered particular interest due to their intrinsic biocompatibility, mechanical compliance, and tunable optical properties, positioning them as promising interfaces for light delivery, biosensing, optogenetic stimulation, and phototherapeutic interventions^25-30^. Recent advances in hydrogel fiber fabrication have enabled precise control over core-cladding architectures, thereby enhancing their optical efficiency and improving functional compatibility with biological tissue environments^25^. However, these hydrogel-based waveguides still face critical challenges, including relatively high optical attenuation, limited refractive index contrast, and structural instability, which restrict their ability to efficiently transmit and preserve high-dimensional optical fields^31^. As a result, the development of hydrogel-based MMF systems capable of delivering high-resolution, high-fidelity images remains an unresolved challenge.

In this study, we introduce HYFEN, a hydrogel-based endomicroscopic MMF imaging platform for flexible, subcellular, and biocompatible fluorescence microscopy. Specifically, HYFEN leverages the unique material and optical properties of hydrogel fibers, in conjunction with adaptive optics and pixel-wise image enhancement, to overcome persistent challenges associated with silica-based MMFs, including mode scrambling, limited field of view (FOV), and mechanical rigidity. This first-of-its-kind hydrogel-based MMF platform enables precise mode-threading, rapid diffraction-limited focusing at kilohertz speeds, and high-fidelity fluorescent signal acquisition with subcellular-scale resolution. Notably, the hydrogel-based approach accommodates substantially enhanced fiber dimensions and bending curvatures that are otherwise unachievable with conventional MMFs, facilitating an extended imaging area without compromising resolution or mechanical integrity. Harnessing advances in materials, optics, and computation, we anticipate that HYFEN will empower minimally invasive imaging and biointerfacing, transforming endoscopic technologies.

## RESULT

### HYFEN: principle and framework

Conventional hydrogel fibers have been explored in various ways due to their mechanical flexibility, biocompatibility, and tunable optical properties (**Supplementary Tables 1 and 2**). In this work, HYFEN leverages custom-fabricated hydrogels, transmission matrix (TM) measurements, adaptive wavefront manipulation, and pixel-wise image enhancement. These strategies collectively enable precise mode threading, effective focusing, and high-quality acquisition of fluorescent signals through multimode hydrogel fibers. It is worth noting that several methods for fabricating hydrogel fibers have been proposed, including tube molding^25,26^, wet spinning^32^, 3D printing^33^, and microfluidic spinning^34^. Here, we selected tube molding for HYFEN due to its simplicity, reproducibility, and precise diameter control (**Methods**). Specifically, polyethylene glycol diacrylate (PEGDA) was used as the fiber core material, polymerized within a microfluidic tube mold through ultraviolet curing. Subsequently, a calcium-alginate hydrogel cladding was formed by immersing the cured PEGDA core into sodium alginate and calcium chloride (CaCl□) solutions^25^ (**Fig. 1a-c**). Alginate was selected as the cladding material due to its low refractive index (1.331), which yields a numerical aperture (NA) greater than 0.48 (**Supplementary Fig. 1, Supplementary Note 1**), thereby enhancing optical confinement and minimizing leakage losses (**Fig. 1d, Supplementary Fig. 2**).

**Figure 1.**
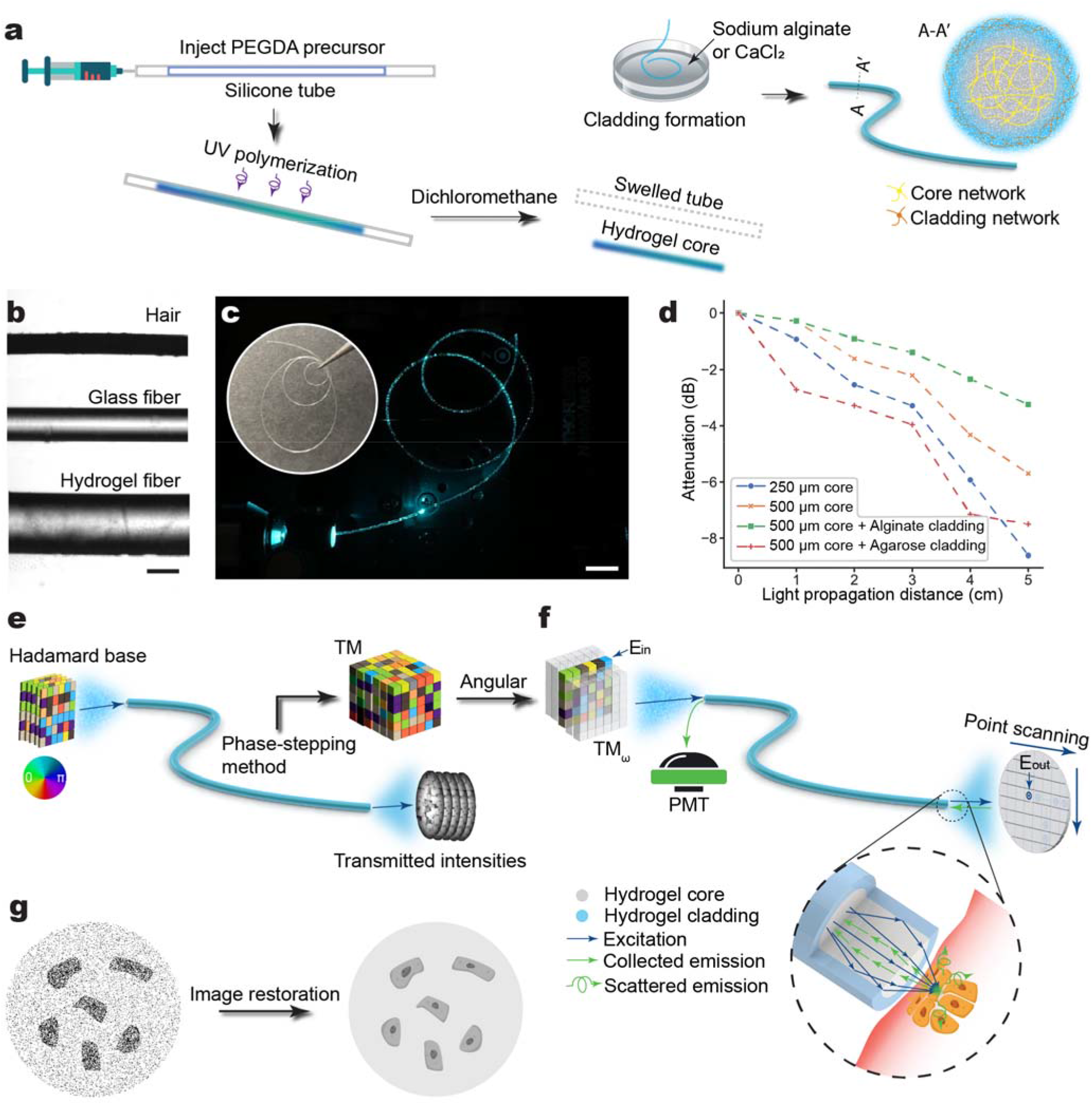
Principle of HYFEN. **(a)** Schematic of the fabrication process for the PEGDA/Ca^2^□alginate hydrogel fiber. **(b)** Comparative images of a human hair, conventional multimode glass fiber (125□µm in diameter), and hydrogel fiber (250□µm in diameter) captured with wide-field microscopy using a 10× objective lens. **(c)** Delivery of 488-nm laser light through a 20-cm deformed hydrogel fiber visualized with a 20× objective lens. Inset: the photograph of the same fiber without light transmission. **(d)** Propagation loss of pre-swollen hydrogel fibers with varying PEGDA core and cladding diameters and settings, quantified via the cutback method^62^. **(e)** Transmission matrix (TM) calibration via phase-stepping. Phase shifts from 0 to 2π are coded with gradient colors. **(f)** Imaging procedures, containing the formation of scanning focal spots via TM inversion and the capture of fluorescence signals through the same hydrogel fiber (inset). **(g)** Schematic of the fluorescence signal extraction and denoising process for biological specimens imaged with HYFEN. Scale bars: 100□µm (b), 10□mm (c).

HYFEN utilizes raster scanning facilitated by computational wavefront shaping. Specifically, we characterized fiber transmission using a reference-free, interference-based TM measurement setup employing a digital micromirror device (DMD)^35^ (**Supplementary Fig. 3**). Hadamard-encoded illumination patterns were introduced at the proximal end of the fiber, and the resulting distal-end speckle patterns were recorded via a CMOS camera and used to compute the TM (**Fig. 1e, Supplementary Fig. 4**). For fast focal spot generation, the modulation and computational rendering were simplified by converting complex phase encoding to angular format (**Supplementary Note 2**). We customized GPU-accelerated CUDA programming for efficient binary pattern encoding on the DMD, achieving a runtime of ∼17.3 seconds for a 256-pixel × 256-pixel image, at least 10 times faster than existing frameworks **(Supplementary Fig. 5**).

For distal-end fluorescence imaging, emitted signals were subsequently collected through the same hydrogel fiber, detected by a photomultiplier tube (PMT), and reconstructed into spatially resolved images using coordinates defined by prior TM measurements (**Fig. 1f**). Notably, single-pixel PMT detectors typically exhibit low quantum efficiency (<39%)^36^, which is often compensated for by increasing laser power^37^. Furthermore, MMF imaging remains susceptible to high background noise, as the power ratio of a single focal spot is physically below 100%^15,38^. Both factors potentially cause photodamage and reduced acquisition rates. Here, HYFEN integrates a lab-developed physical noise model and a microlocal pixel-wise shrinkage algorithm, leveraging shearlet-based optimal sparse representation for robust signal extraction from noisy data^39^. Therefore, the system circumvents these limitations by effectively utilizing PMT detection at lower laser intensities and shorter pixel dwell times (∼50 μs), allowing for faster acquisition without compromising sample health and underlying signal details (**Fig. 1g, Methods**).

### Characterization of HYFEN

The imaging performance of optical fibers depends on several factors, such as focusing uniformity, achievable spatial resolution, and the enhancement factor, which determines the signal-to-noise ratio (SNR) of reconstructed fluorescence images^38^. To characterize the performance of HYFEN, we initially compared tightly focused spots generated by conventional silica glass fibers and hydrogel fibers. The TM measurements and adaptive wavefront shaping achieved focusing of otherwise speckled patterns across defined spatial coordinates (**Fig. 2a, b**). As measured, the hydrogel fiber exhibited a comparable focusing pattern (theoretical resolution after deconvolution was 1.08 µm) due to its relatively high NA (>0.48) compared to conventional glass fiber, including those with higher NA (**Fig. 2c, d, Supplementary Fig. 6**). Moreover, despite the increased optical attenuation in hydrogel fibers due to surface irregularities at the core-cladding interface and refractive index variations from inhomogeneous cross-linking, HYFEN achieves an enhancement factor comparable to that of glass fibers, delivering a 2 to 3 orders of magnitude improvement across an expanded FOV. Due to higher optical attenuation and larger mode numbers resulting from increased fiber diameters, HYFEN exhibits a disparity in the enhancement factor compared to the conventional system (**Fig. 2e, f**). This enhancement factor maintains essential transmission efficiency, decreasing slightly over distances of several centimeters (**Fig. 2g**). Notably, the enhancement factor remains steady over extended time courses (tens of minutes), demonstrating robustness despite the soft, flexible nature of hydrogel fibers, which would typically lead to rapid performance deterioration compared to rigid glass fibers (**Fig. 2g, Supplementary Note 3, Supplementary Fig. 7**). In addition, we demonstrated the spatial accuracy and scanning capability of HYFEN by programming and sequentially constructing wide-field images (250 × 250 µm) at the fiber output facet, achieving an over 2-fold enhancement in the FOV compared to standard glass fibers (**Fig. 2h**). Additionally, we employed experimentally measured illumination foci to synthesize and calibrate imaging performance based on a standard 1951 USAF resolution test target (**Fig. 2i, Supplementary Note 4**). HYFEN effectively resolved microscopic features, including 228 line pairs/mm across the FOV of 250 × 250 µm, which alternatively required multiple scanning iterations with conventional glass fibers (**Fig. 2j, k**).

**Figure 2.**
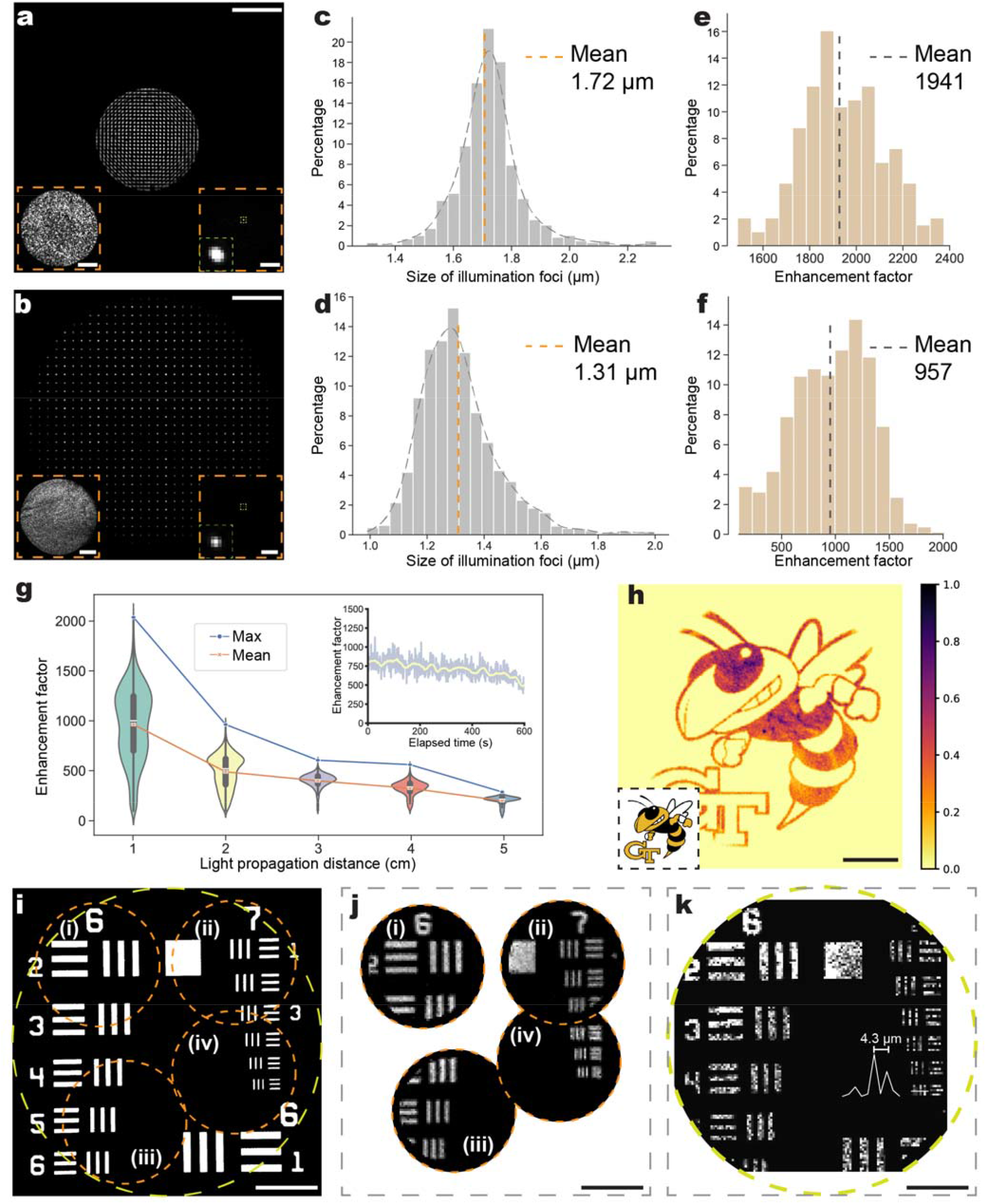
Optical characterization of the HYFEN platform. **(a, b)** Cumulative focal maps at the distal facets of a multimode glass fiber (105□µm in diameter) (a) and a hydrogel fiber (250□µm in diameter) (b), demonstrating uniform focusing at the output. Insets show output speckles without (left) and foci with (right) phase modulation and corresponding zoomed-in focal image (green). **(c, d)** Spatial profile distributions of 400 illumination foci at the distal facets of the glass (c) and hydrogel (d) fibers. **(e, f)** Distribution of the enhancement factor (*S*_peak_/*S*_average_) corresponding to the same 400 illumination foci at the distal end of the glass (e) and hydrogel (f) fibers. **(g)** Enhancement factor as a function of propagation distance in hydrogel fibers. Inset: temporal stability of the enhancement factor. **(h)** Buzz, the Georgia Tech mascot, formed with cumulative foci through a hydrogel fiber. Inset: the original input image. **(i)** 1951 USAF resolution test target over a 250□×□250□µm FOV. (**j, k**) Synthetic MMF images, formed by convolving the experimental foci of multimode glass (j) and hydrogel (k) fibers with the corresponding regions in the test target (i). The cross-sectional profile in (k) indicates resolved element 6, group 7 of the test target. The characterizations were performed with a fiber length *L* = 1 cm (a-f). Scale bars: 50 µm (a, b, h-k), 25 µm (a, b inset).

### Fluorescence microscopy using HYFEN

Next, we imaged fluorescent microspheres with diameters of 4 µm and 15 µm using HYFEN (**Fig. 3a, b**). To minimize photobleaching, we applied a continuous-wave 488-nm laser at a maximum power of 200 µW per focal spot, covering an imaging area with a diameter of 250 µm. As assessed, the pixel-wise image processing of HYFEN demonstrated wide-field robustness for single-pixel PMT sensors by effectively reducing background noise and improving structural similarity to the reference images (**Supplementary Table 3**). We then utilized HYFEN to image the nuclei of cultured HeLa cells labeled with Syto16, which captured the subcellular mitotic phase (<10 μm) in dividing cells with higher structural clarity (**Fig. 3c**).

**Figure 3.**
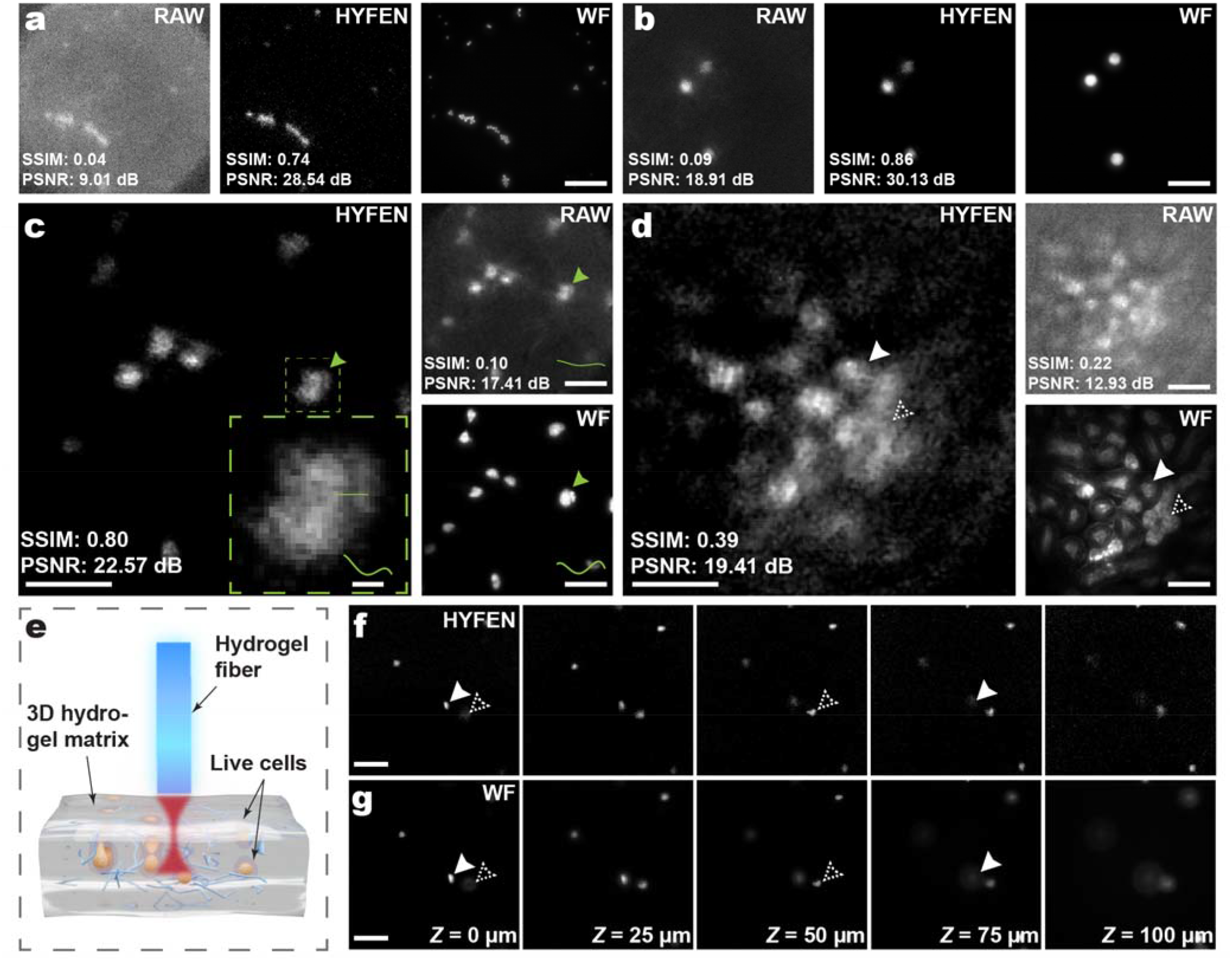
Fluorescence microscopy using HYFEN. **(a, b)** Raw hydrogel fiber, HYFEN, and wide-field (WF, ground truth) images of fluorescent microspheres with diameters of 4□µm (a) and 15□µm (b). **(c)** Raw hydrogel fiber, HYFEN, and wide-field images of Syto16-labeled nuclei in HeLa cells. Inset: zoomed-in image of the dashed boxed region in the HYFEN image, resolving microscopic sub-cellular features (arrow-pointed) in the cross-section profiles. **(d)** Raw hydrogel fiber, HYFEN, and wide-field images of a WGA-labeled mouse kidney tissue section. The dashed and solid arrows indicate the resolution of glomeruli and convoluted tubules, respectively. **(e)** Schematic of HYFEN imaging of live cells embedded in a 3D hydrogel matrix. **(f, g)** HYFEN and wide-field images of live cells taken at an axial step size of 25 µm across a depth range of 100□µm. The dashed and solid arrows mark the cells acquired at different axial positions. Imaging was performed with a fiber length *L* = 2 cm. Scale bars: 50□µm (a-d, f, g), 10□µm (c, inset).

Furthermore, we extended HYFEN to image a cryostat section of a mouse kidney stained with Alexa Fluor 488-conjugated WGA (wheat germ agglutinin) lectin, which targets N-acetylglucosamine sugar and sialic acid residues. HYFEN resolved fine single-cell details in the tissue samples, achieving a rapid scan time of 3.3 seconds for an area of 250 × 250 µm with an extensive focal range of 18.9 µm, capturing the entire 16-µm-thick tissue section. In particular, HYFEN allows for discerning high-contrast patterns of various renal cellular morphology, effectively distinguishing glomerular and tubular components within the mouse kidney (**Fig. 3d**). In addition, to test optical sectioning, we imaged live, 3D-cultured HeLa cells distributed within a hydrogel matrix (**Fig. 3e**). HYFEN acquired precise 3D single-cell positions across a 100-μm depth range and resolved nearby cells as close as ∼7 µm and 50 µm in the lateral and axial dimensions, respectively, in agreement with the results of the same region obtained by a 10× microscope objective lens (**Fig. 3f, g, Supplementary Fig. 8**).

### High-resolution imaging with enhanced fiber flexibility and extended FOV

Conventional MMFs encounter a typical trade-off between fiber flexibility and diameter, which limits their biointerfacing ability and achievable FOV. Larger fiber diameters increase rigidity, often resulting in brittle fibers that are prone to breakage, thereby heightening the risk of tissue damage during imaging procedures. In contrast, soft hydrogel fibers maintain flexibility and allow for an extended usable FOV and robustness under tighter bending (i.e., small radii of curvature). Here, we implemented HYFEN with a thicker hydrogel fiber exceeding 500 µm in diameter, which correspondingly enabled an effective FOV over 400□µm in diameter, a dimension that becomes impractical with standard glass-based MMFs due to their rigidity and fragility (**Fig. 4a**). We further demonstrated that by combining sample scanning, HYFEN can be extended to screen large cell populations with subcellular resolution at a steady throughput of 100-1,000 cells per second (**Fig. 4b, c**). Notably, despite the quadratic increase in supported fiber modes associated with the larger fiber core, HYFEN maintained robust imaging performance, generating consistent segmentation and profiling of cell radii between 10 µm and 40 µm and populational structural similarity (>70%), in comparison with standard wide-field microscopy (**Fig. 4d, e, Supplementary Fig. 9, Methods**).

**Figure 4.**
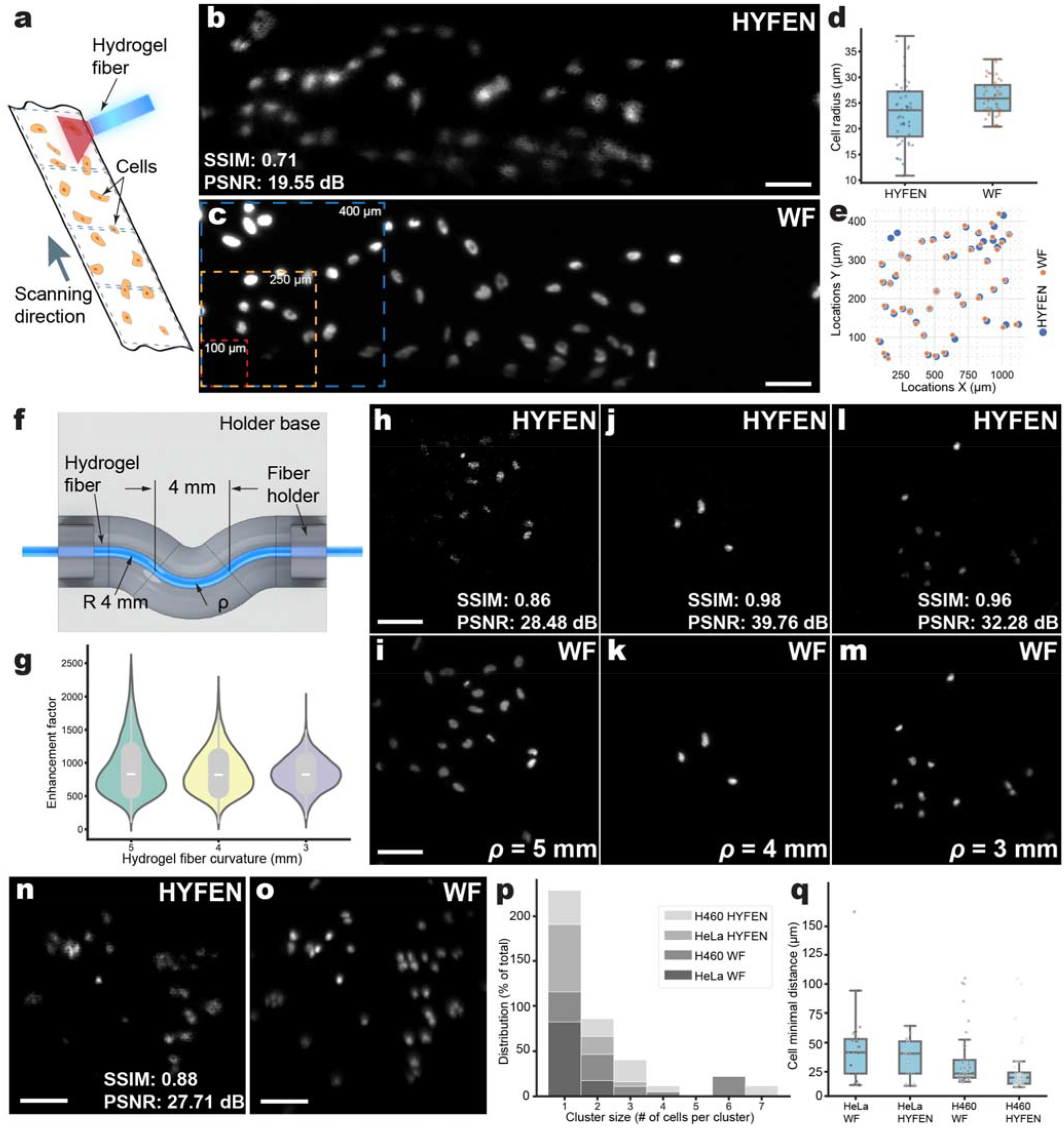
High-resolution HYFEN imaging with enhanced fiber flexibility and extended FOV. **(a)** Schematic of HYFEN imaging with sample scanning to achieve an extended FOV. **(b, c)** HYFEN (b) and wide-field (c) fluorescence imaging of the nuclei in HeLa cells over a FOV of 410□µm (width) × 1467□µm (length). Red, yellow, and blue boxes indicate regions of 100□×□100□µm, 250□×□250□µm, and 400□×□400□µm, respectively. **(d, e)** Quantification of nuclear radii (d) and spatial distribution (e) from HYFEN and wide-field images. **(f)** Schematic of a hydrogel fiber placed in a 3D-printed holder configured for tight bending. **(g)** Enhancement factors of HYFEN under different radii of curvature (ρ = 5, 4, and 3 mm). **(h-m)** Comparison of HYFEN (h, j, l) and wide-field (i, k, m) images of HeLa cells at bending curvatures of ρ = 5 mm, 4 mm, and 3 mm, respectively. **(n, o)** HYFEN (n) and wide-field (o) images of the nuclei in H460 cells. **(p)** Cluster size analysis of HeLa and H460 nuclei using DBSCAN, showing the distribution of cell types grouped by cluster size. **(q)** Distribution of nearest-neighbor distance, quantifying intercellular spacing of HeLa and H460 nuclei under HYFEN and wide-field imaging. Imaging was performed with a fiber length *L* = 2 cm. Scale bars: 100□µm.

Lastly, to validate flexibility, we sequentially positioned a 20 mm-long hydrogel fiber into custom-fabricated microchannels designed with varying radii (ρ = 5, 4, and 3 mm), which correspondingly indicated tighter bending curvatures (**Fig. 4f, Supplementary Fig. 10**). Here, HYFEN exhibited consistent enhancement factors exceeding 750× for all curvatures, without noticeable light leakage (**Fig. 4g**), and preserved the high-resolution imaging capabilities of HeLa cell nuclei (**Fig. 4h-m**). With a pre-measured TM database, HYFEN demonstrated resistance to bending and temperature perturbations (**Supplementary Fig. 11, Supplementary Movie 1**). Furthermore, using curved hydrogel fibers (ρ = 5□mm), we imaged H460 lung cancer cells, which characteristically form clusters rather than disperse as single cells^40,41^ (**Fig. 4n, o**). We conducted cluster size analysis and quantified that at least 62% of H460 cells were part of clusters containing two or more cells, in contrast to approximately 25% of HeLa cells (**Fig. 4p, Methods**). Furthermore, image analysis of nearest-neighbor distances revealed reduced intercellular spacing for H460 cells (75% of H460 cells below 18□µm) compared to HeLa cells (27□µm), consistent with the measurements obtained by standard wide-field microscopy (**Fig. 4q**). Collectively, these results validated the capability of HYFEN to achieve high-resolution fluorescence imaging under enhanced fiber dimensions and flexibility, conditions that otherwise unachievable with conventional MMFs.

## DISCUSSION

MMFs offer minimally invasive, high-resolution imaging through ultrathin probes, enhancing diagnostic precision and enabling real-time monitoring in delicate anatomical regions. In this work, we present HYFEN, a hydrogel-based MMF imaging platform that exploits the intrinsic biocompatibility and flexibility of hydrogels while addressing previous challenges, including optical attenuation, mode scrambling, and material stiffness. HYFEN leverages innovative soft materials, adaptive optics, and pixel-wise image enhancement to achieve subcellular resolution, exceptional mechanical flexibility, and an extended FOV compared to conventional silica-based MMFs. The robustness of the method has been demonstrated under various tissue samples and environmental settings (**Supplementary Figs. 12-14, Supplementary Notes 5-7**).

The methodology can be further advanced with novel materials^42-44^, optical configurations^45,46^, and computational frameworks^47-50^. Like conventional MMF systems, HYFEN is susceptible to bendinginduced mode scrambling, which can be mitigated by constructing a TM database with full-vector modulation to ensure robust imaging performance under complex biological conditions^19^. Environmental perturbations, including temperature fluctuations, airflow, and DMD thermal load, contribute to a decline in the enhancement factor (**Supplementary Fig. 15**). This can be alleviated by incorporating a thermoelectric cooler (TEC) and heat sink for DMD temperature stabilization^51^ and minimizing the optical path length to reduce airflow effects. In addition, the structural stability of the hydrogel optical fiber can be further enhanced by using core-cladding materials that form robust covalent bonds, mitigating temporary detachment observed during swelling (**Supplementary Movie 2**)^52,53^. Future improvements will involve post-polishing to achieve a smoother end face, enhancing the imaging quality of HYFEN.

We anticipate the accessible platform to enable versatile integration with a broad range of hydrogelmediated biomedical technologies, including optogenetics^28,54^, tissue engineering constructs^55,56^, and implantable medical devices^57,58^. HYFEN will serve as a powerful paradigm for biointerfacing and medical imaging applications, enabling the exploration of complex biomedical systems beyond conventional material, optical, and computational limitations.

## METHODS

### Experimental system

We used wavefront-shaping to modulate the phase input at the proximal end of the hydrogel fiber and generated raster-scanning foci at the distal end of the fiber with micron-scale resolution (**Supplementary Fig. 3, Supplementary Table 4**). A continuous-wave laser with a wavelength of 488 nm was used as the excitation source, and the laser beam was expanded using a 4-*f* system. The expanded beam was then incident on DMD at 12°, and the modulated first-order diffraction beam was selected by an iris at the Fourier plane, which was conjugated to the focus plane of the objective lens and the proximal end of the fiber. The speckled outputs of the fiber were collected by another objective lens and tube lens, then detected by a camera. Off-axis binary amplitude holography, based on the Lee method, was used to encode the target phases using a DMD^59,60^. With the Hadamard-matrix-based phase sequences generated by the DMD, the camera can record speckles at a rate of up to 1,000 frames per second, triggered by the DMD. After the measurement of the TM of the fiber, the scanning foci were generated, and the fluorescence signals from the samples were transmitted back through the fiber and filtered with a dichroic mirror and a bandpass emission filter, then detected by a photon-sensitive PMT. PMT signals were acquired and synchronized with the DMD using a data acquisition card and controlled via its open-source API (NIDAQmx-python, National Instruments). We used a customized phase width modulator (PWM) to multiply the trigger signals (×10) from the DMD and averaged the detected signals from the PMT to further improve the SNR (**Supplementary Note 4**). The refresh rate of the DMD for imaging was 20,000 Hz, and the frame rate per reconstructed fluorescence image with a pixel size of 256 × 256 was 0.31 frames per second.

### Fabrication of the hydrogel fiber

Biocompatible microfluidic tubes with an inner diameter of 250 µm or 510 µm were used as molds for the hydrogel cores. For core preparation, an 80% w/v PEGDA and 5% w/v 2-hydroxy-2-methylpropiophenone solution in deionized water (DW) was thoroughly mixed using a vortex mixer as the precursor solution. The solution was injected into the tube mold and exposed to UV light for 15 minutes for photo-crosslinking. The core was then demolded from the tube, which had been treated with dichloromethane. 3% w/v sodium alginate solution was prepared by dissolving 3.0 g of sodium alginate in 100 mL of DW within a 250-mL Erlenmeyer flask. A magnetic stir bar was added, and the solution was stirred at 600 revolutions per minute (rpm) for approximately 1 hour, or until it was fully dissolved. The hydrogel fiber cores were immersed in a sodium alginate solution for 1 minute, followed by a 3-minute drying step. Subsequently, the fibers were submerged in a 100 mmol/L CaCl□ solution for 1 minute to induce Ca^2+^-alginate cladding formation. The cladding formation process could be repeated for multiple layers. The hydrogel fiber was cut to the target length with a razor blade and stored in DW before imaging. All the category information on necessary materials and equipment can be found in **Supplementary Table 5** and **Supplementary Note 8**.

### Fluorescence beads slide

The imaging quality of the hydrogel fiber was first evaluated with 4 µm and 15 µm beads. The beads are used at ratios of 1:10, 1:20, 1:50, or 1:100 to achieve varying concentrations from dense to sparse. The coverslip was first depolarized using an electric gun, which was placed on an aluminum foil in a clear surrounding area, and the charge was applied for 30 to 60 seconds. A 20 µL aliquot of the diluted bead solution was then pipetted onto the charged coverslip and spread evenly across its entire surface. The coverslip was subsequently placed in a closed box protected from light and left to dry for 6 hours. 20 µL glycerol solution was pipetted onto a glass slide. The dried coverslip was then carefully flipped, ensuring the side containing the beads was placed onto the glycerol-coated slide and evenly spread. The edges of the coverslip were sealed using transparent nail polish, which was left to dry for 15 to 30 minutes before imaging.

### HeLa and H460 cell culture

HeLa cells are derived from cervical carcinoma, and H460 cells are derived from lung adenocarcinoma. Both cell lines were cultured in cell culture medium (For HeLa cells, Dulbecco’s Modified Eagle Medium (DMEM); for H460 cells, Roswell Park Memorial Institute (RPMI)) supplemented with 10% fetal bovine serum (FBS) and 1% Penicillin-Streptomycin (Pen-Strep) at 37□°C under a 5% CO_2_ atmos-phere.

### Nuclei labeled with fixed HeLa and H460 cells

A single coverslip was placed in a 35-mm culture dish containing gelatin and treated for 30 minutes. Subsequently, 2 mL DMEM was added, followed by the addition of 200 µL of HeLa or H460 cell suspension. The cells were allowed to grow and attach to the coverslip for 2–3 days. After incubation, the DMEM was removed from the culture dish. The labeling reagent was prepared with a concentration of 5 µM in a plastic tube by mixing 2 mL of FluoroBrite DMEM with 10 µL of Syto16 solution. The prepared labeling solution was prewarmed to 37 °C and then added to the culture dish. The cells were incubated for 1 hour at 37°C under a CO_2_ atmosphere. After labeling, the solution was removed, and the cells were washed three times with phosphate-buffered saline (PBS). For fixation, cells were treated with 4% paraformaldehyde (PFA) for 12 minutes inside the incubator. After fixation, the PFA was removed, and the cells were washed with PBS. The coverslip was carefully removed from the culture dish, and 25 µL of liquid antifade mountant was applied to a glass slide to minimize photobleaching. The coverslip was then placed onto the glass slide and secured with nail enamel, which was left to dry for 15 to 30 minutes before imaging.

### Nuclei labeled 3D live HeLa cells in the hydrogel matrix

10 mL of 2% w/v sodium alginate in PBS and 10 mL of 100 mmol/L CaCl□ solution in PBS were prepared and prewarmed to 37°C before use. 2 mL of the CaCl□ solution was added to a 35-mm culture dish. (The precursor of the alginate hydrogel). The HeLa cell suspension with 2 mL DMEM was prepared in a 15 mL plastic tube and centrifuged for 5 minutes at 300 relative centrifugal force (RCF). The cells were resuspended in 1 mL DMEM. 1 mL sodium alginate solution and 500 µL cell medium were added to a 5 mL plastic tube and mixed. The cell and sodium alginate mixture was then pipetted into the CaCl□ dish to form the hydrogel spheres. The CaCl□ solution was removed, and a 5 µM Syto16 solution in FluoroBrite DMEM was prepared and added to the hydrogel dish for labeling. The hydrogel dish was incubated for 1 hour at 37°C under a CO_2_ atmosphere. After labeling, the labeling solution was removed, and the cells were washed three times with PBS. The hydrogel spheres were then removed from the dish and placed on a glass slide, which was then covered with a coverslip. Nail enamel was used to seal the coverslip and left to dry for 15 to 30 minutes before imaging.

### HYFEN image restoration

Before proceeding with sample imaging, the fluorescence tape was placed in front of the hydrogel fiber, and the signals were captured as the “Offset” input to the HYFEN system. The emission wavelength, minimal resolution, and system magnification were set to 515 nm, 1.2 µm, and 10×. Multiscale Wiener filtering and a pre-calculated shearlet system were employed to achieve maximum performance^39^. The average processing speed per image (256 × 256 pixels) was 2.38 seconds.

### CellProfiler for cell segmentation

We used the IdentifyPrimaryObjects function in CellProfiler for both HYFEN and wide-field imaging cell segmentation (**Supplementary Fig. 16**). To cover weak signals, the typical diameters of detected objects were set from 7 to 42 pixels. Objects outside this diameter range and those touching the image borders were discarded. The global threshold strategy was adopted, with minimum cross-entropy as the thresholding method. After segmentation, the major and minor axis lengths, as well as the center location of each cell, were exported for further statistical analysis and visualization.

### Distribution measurement of the cluster sizes

Following cell segmentation using CellProfiler, the exported cell coordinates were converted from pixel units to physical distances in microns. We applied Density-Based Spatial Clustering of Applications with Noise (DBSCAN)^61^ to determine cluster sizes for the HYFEN and wide-field imaging datasets of H460 and HeLa cells. The maximum distance between two cells for them to be considered neighbors was set to 16.5 µm for optimum clustering, and the minimum number of cells (or total weight) required for a core point was set to 2. Isolated cells (classified as noise) were treated as single-cell clusters. The resulting cluster sizes were visualized as percentage distributions.

## DATA AVAILABILITY

All data needed to evaluate the conclusions in the paper are present in the paper and/or the Supplementary Materials. Additional data is publicly available here: https://doi.org/10.5281/zenodo.17195639.

## CODE AVAILABILITY

The codes have been written in Python and tested with version 3.11.5. To run the codes, unzip the compressed folder and follow the instructions in the file readme.md. The most updated version of the testing codes can be found at: https://github.com/ShuJiaLab/HYFEN.

## ACKNOWLEDGEMENTS

We acknowledge the support of the National Institutes of Health grants R35GM124846 (to S.J.), the National Science Foundation grants 2145235 and 2503686 (to S.J.), Georgia Institute of Technology Faculty Startup Fund (to S.J.), the National Science Foundation Graduate Research Fellowship DGE-2039655 (to C.Z.), and Career Award at the Scientific Interface from the Burroughs Wellcome Fund (to J.E.M.). We thank Wenhao Liu of the Georgia Institute of Technology and Shengfu Cheng of the Hong Kong Polytechnic University for their helpful advice on this work, and Didun Oladeji of the Georgia Institute of Technology for her assistance with refractive index measurements.

## AUTHOR CONTRIBUTIONS STATEMENT

P.C. and S.J. conceived and designed the project. P.C., K.H., Z.G., H.X., C.Z., and M.T.C. contributed to the construction of the optical system and fabrication of the hydrogel fibers. M.S. and A.H.K prepared and provided live cell samples. C.M.D. and J.E.M. prepared and provided mouse brain samples. P.C. and Z.L. performed the image post-processing. P.C. performed imaging experiments. S.J. supervised the overall project. P.C. and S.J. wrote the manuscript with input from all authors.

## REFERENCES

1 Tsuboi, M. et al. Optical coherence tomography in the diagnosis of bronchial lesions. Lung cancer 49, 387–394 (2005).

2 Subramanian, V. & Ragunath, K. Advanced endoscopic imaging: a review of commercially available technologies. Clinical Gastroenterology and Hepatology 12, 368-376. e361 (2014).

3 Gora, M. J., Suter, M. J., Tearney, G. J. & Li, X. Endoscopic optical coherence tomography: technologies and clinical applications. Biomed. Opt. Express, BOE 8, 2405–2444 (2017).

4 Orth, A., Ploschner, M., Wilson, E. R., Maksymov, I. S. & Gibson, B. C. Optical fiber bundles: Ultra-slim light field imaging probes. Science Advances 5, eaav1555 (2019). 10.1126/sciadv.aav1555

5 Urner, T. M., Inman, A., Lapid, B. & Jia, S. Three-dimensional light-field microendoscopy with a GRIN lens array. Biomed. Opt. Express, BOE 13, 590–607 (2022). 10.1364/BOE.447578

6 Cao, H., Čižmár, T., Turtaev, S., Tyc, T. & Rotter, S. Controlling light propagation in multimode fibers for imaging, spectroscopy, and beyond. Advances in Optics and Photonics 15, 524–612 (2023).

7 Psaltis, D. & Moser, C. Imaging with multimode fibers. Optics and Photonics News 27, 24–31 (2016).

8 Mahalati, R. N., Gu, R. Y. & Kahn, J. M. Resolution limits for imaging through multi-mode fiber. Optics express 21, 1656–1668 (2013).

9 Čižmár, T. & Dholakia, K. Exploiting multimode waveguides for pure fibre-based imaging. Nat Commun 3, 1027 (2012). 10.1038/ncomms2024

10 Plöschner, M., Tyc, T. & Čižmár, T. Seeing through chaos in multimode fibres. Nature Photonics 9, 529–535 (2015). 10.1038/nphoton.2015.112

11 Plöschner, M. et al. Multimode fibre: Light-sheet microscopy at the tip of a needle. Sci Rep 5, 18050 (2015). 10.1038/srep18050

12 Morales-Delgado, E. E., Psaltis, D. & Moser, C. Two-photon imaging through a multimode fiber. Opt. Express, OE 23, 32158–32170 (2015). 10.1364/OE.23.032158

13 Caravaca-Aguirre, A. M. & Piestun, R. Single multimode fiber endoscope. Opt. Express, OE 25, 1656–1665 (2017). 10.1364/OE.25.001656

14 Ohayon, S., Caravaca-Aguirre, A., Piestun, R. & DiCarlo, J. J. Minimally invasive multimode optical fiber microendoscope for deep brain fluorescence imaging. Biomed. Opt. Express, BOE 9, 1492–1509 (2018). 10.1364/BOE.9.001492

15 Turtaev, S. et al. High-fidelity multimode fibre-based endoscopy for deep brain in vivo imaging. Light Sci Appl 7, 92 (2018). 10.1038/s41377-018-0094-x

16 Vasquez-Lopez, S. A. et al. Subcellular spatial resolution achieved for deep-brain imaging in vivo using a minimally invasive multimode fiber. Light Sci Appl 7, 110 (2018). 10.1038/s41377-018-0111-0

17 Stibůrek, M. et al. 110 μm thin endo-microscope for deep-brain in vivo observations of neuronal connectivity, activity and blood flow dynamics. Nat Commun 14, 1897 (2023). 10.1038/s41467-023-36889-z

18 Stellinga, D. et al. Time-of-flight 3D imaging through multimode optical fibers. Science 374, 1395–1399 (2021). 10.1126/science.abl3771

19 Wen, Z. et al. Single multimode fibre for in vivo light-field-encoded endoscopic imaging. Nature Photonics 17, 679–687 (2023). 10.1038/s41566-023-01240-x

20 Cifuentes, A. et al. Polarization-resolved second-harmonic generation imaging through a multimode fiber. Optica, OPTICA 8, 1065–1074 (2021).

21 Liu, X., Liu, J., Lin, S. & Zhao, X. Hydrogel machines. Materials Today 36, 102–124 (2020).

22 Parker, S. T. et al. Biocompatible silk printed optical waveguides. Advanced Materials 21 (2009).

23 Orelma, H. et al. Optical cellulose fiber made from regenerated cellulose and cellulose acetate for water sensor applications. Cellulose 27, 1543–1553 (2020).

24 Xin, H., Li, Y., Liu, X. & Li, B. Escherichia coli-based biophotonic waveguides. Nano Lett. 13, 3408–3413 (2013).

25 Choi, M., Humar, M., Kim, S. & Yun, S.-H. Step-Index Optical Fiber Made of Biocompatible Hydrogels. Advanced Materials 27, 4081–4086 (2015). 10.1002/adma.201501603

26 Yetisen, A. K. et al. Glucose□sensitive hydrogel optical fibers functionalized with phenylboronic acid. Advanced Materials (Deerfield Beach, Fla.) 29, 1606380 (2017). 10.1002/adma.201606380

27 Jiang, N. et al. Functionalized flexible soft polymer optical fibers for laser photomedicine. Advanced Optical Materials 6, 1701118 (2018). 10.1002/adom.201701118

28 Liu, X. et al. Fatigue-resistant hydrogel optical fibers enable peripheral nerve optogenetics during locomotion. Nat Methods, 1–8 (2023). 10.1038/s41592-023-02020-9

29 Sadeque, M. S. B. et al. Hydrogel-integrated optical fiber sensors and their applications: a comprehensive review. J. Mater. Chem. C 11, 9383–9424 (2023). 10.1039/D3TC01206A

30 Zhang, Z. et al. Dual-function wearable hydrogel optical fiber for monitoring posture and sweat pH. ACS sensors 9, 3413–3422 (2024).

31 Yew, P. Y. M. et al. Hydrogel for light delivery in biomedical applications. Bioactive Materials 37, 407–423 (2024).

32 Chen, G. et al. Integrated dynamic wet spinning of core-sheath hydrogel fibers for optical-to-brain/tissue communications. National Science Review 8, nwaa209 (2021). 10.1093/nsr/nwaa209

33 Lorang, D. J. et al. Photocurable Liquid Core–Fugitive Shell Printing of Optical Waveguides. Advanced Materials 23, 5055–5058 (2011). 10.1002/adma.201102411

34 Daniele, M. A. et al. Rapid and Continuous Hydrodynamically Controlled Fabrication of Biohybrid Microfibers. Advanced Functional Materials 23, 698–704 (2013). 10.1002/adfm.201202258

35 Popoff, S. M. et al. Measuring the Transmission Matrix in Optics: An Approach to the Study and Control of Light Propagation in Disordered Media. Physical Review Letters 104, 100601 (2010). 10.1103/PhysRevLett.104.100601

36 Hamamatsu Photonics, K. Photomultiplier tubes: Basics and applications. Edition 3a 310 (2007).

37 Jonkman, J., Brown, C. M., Wright, G. D., Anderson, K. I. & North, A. J. Tutorial: guidance for quantitative confocal microscopy. Nature protocols 15, 1585–1611 (2020).

38 Gomes, A. D., Turtaev, S., Du, Y. & Čižmár, T. Near perfect focusing through multimode fibres. Opt. Express, OE 30, 10645–10663 (2022). 10.1364/OE.452145

39 Mandracchia, B. et al. Optimal sparsity allows reliable system-aware restoration of fluorescence microscopy images. Science Advances 9, eadg9245 (2023).

40 Zhang, X., Xu, L.-h. & Yu, Q. Cell aggregation induces phosphorylation of PECAM-1 and Pyk2 and promotes tumor cell anchorage-independent growth. Molecular cancer 9, 1–11 (2010).

41 Kwon, S., Yang, W., Moon, D. & Kim, K. S. Comparison of cancer cell elasticity by cell type. Journal of Cancer 11, 5403 (2020).

42 Dong, Y. C., Bouche, M., Uman, S., Burdick, J. A. & Cormode, D. P. Detecting and monitoring hydrogels with medical imaging. ACS biomaterials science & engineering 7, 4027–4047 (2021).

43 Park, G. K. et al. Dual-Channel Fluorescence Imaging of Hydrogel Degradation and Tissue Regeneration in the Brain. Theranostics 9, 4255–4264 (2019). 10.7150/thno.35606

44 Li, X. & Gong, J. P. Design principles for strong and tough hydrogels. Nature Reviews Materials 9, 380–398 (2024).

45 Qiu, T. et al. Spectral-temporal-spatial customization via modulating multimodal nonlinear pulse propagation. Nature Communications 15, 2031 (2024). 10.1038/s41467-024-46244-5

46 Lamb, E. S., Shi, Z., Kremp, T., DiGiovanni, D. J. & Westbrook, P. S. Shape sensing endoscope fiber. Optica 11, 1462–1467 (2024). 10.1364/OPTICA.532250

47 Rahmani, B., Loterie, D., Konstantinou, G., Psaltis, D. & Moser, C. Multimode optical fiber transmission with a deep learning network. Light: science & applications 7, 69 (2018).

48 Liu, Z. et al. All-fiber high-speed image detection enabled by deep learning. Nature communications 13, 1433 (2022).

49 Xue, Y., Yang, Q., Hu, G., Guo, K. & Tian, L. Deep-learning-augmented computational miniature mesoscope. Optica 9, 1009–1021 (2022).

50 Valzania, L. & Gigan, S. Online learning of the transmission matrix of dynamic scattering media. Optica 10, 708–716 (2023). 10.1364/OPTICA.479962

51 Rudolf, B., Du, Y., Turtaev, S., Leite, I. T. & Čižmár, T. Thermal stability of wavefront shaping using a DMD as a spatial light modulator. Opt. Express, OE 29, 41808–41818 (2021). 10.1364/OE.442284

52 Liu, B. et al. Hydrogel Coating Enabling Mechanically Friendly, Step-Index, Functionalized Optical Fiber. Advanced Optical Materials 9, 2101036 (2021). 10.1002/adom.202101036

53 Yuk, H., Zhang, T., Lin, S., Parada, G. A. & Zhao, X. Tough bonding of hydrogels to diverse non-porous surfaces. Nature Mater 15, 190–196 (2016). 10.1038/nmat4463

54 Choi, M. et al. Light-guiding hydrogels for cell-based sensing and optogenetic synthesis in vivo. Nature Photonics 7, 987–994 (2013). 10.1038/nphoton.2013.278

55 Cao, H., Duan, L., Zhang, Y., Cao, J. & Zhang, K. Current hydrogel advances in physicochemical and biological response-driven biomedical application diversity. Signal transduction and targeted therapy 6, 426 (2021).

56 Chen, X. et al. Hydrogel Fibers-Based Biointerfacing. Advanced Materials 37, 2413476 (2025). 10.1002/adma.202413476

57 Ren, H. et al. Injectable, self-healing hydrogel adhesives with firm tissue adhesion and on-demand biodegradation for sutureless wound closure. Science Advances 9, eadh4327 (2023). doi:10.1126/sciadv.adh4327

58 Tagami, T. et al. 3D printing of gummy drug formulations composed of gelatin and an HPMC-based hydrogel for pediatric use. International Journal of Pharmaceutics 594, 120118 (2021).

59 Lee, W.-H. Binary computer-generated holograms. Appl. Opt., AO 18, 3661–3669 (1979). 10.1364/AO.18.003661

60 Conkey, D. B., Caravaca-Aguirre, A. M. & Piestun, R. High-speed scattering medium characterization with application to focusing light through turbid media. Opt. Express, OE 20, 1733–1740 (2012). 10.1364/OE.20.001733

61 Ester, M., Kriegel, H.-P., Sander, J. & Xu, X. in kdd. 226–231.

62 Guo, J. et al. Highly stretchable, strain sensing hydrogel optical fibers. Advanced Materials 28, 10244–10249 (2016). 10.1002/adma.201603160

